# Structural basis for the access and binding of resolvin D1 (RvD1) to formyl peptide receptor 2 (FPR2/ALX), a class A GPCR

**DOI:** 10.1101/2024.09.23.614540

**Authors:** Brian Mun, Peter Obi, Christopher T. Szlenk, Senthil Natesan

## Abstract

Inflammation is essential to the body’s defense against tissue injury and microbial invasion. However, uncontrolled inflammation is highly detrimental and can result in chronic inflammatory diseases such as asthma, cancer, obesity, and diabetes. An increasing body of evidence suggests that specialized pro-resolving lipid mediators (SPMs), such as resolvins, are actively involved in critical cellular events that drive the resolution of inflammation and a return to homeostasis. An imbalance caused by insufficient SPMs can result in the unsuccessful resolution of inflammation. The D-series resolvins (metabolites of docosahexaenoic acid), such as resolvin D1 (RvD1) and resolvin D2 (RvD2), carry out their pro-resolving functions by directly binding to class A G protein-coupled receptors FPR2/ALXR and GPR32, and GPR18, respectively. We recently demonstrated that RvD1 and RvD2 preferentially partition and accumulate at the polar headgroup regions of the membrane. However, the mechanistic detail of how RvD1 gains access to the FPR2 binding site from a surrounding membrane environment remains unknown. In this study, we used classical MD and well-tempered metadynamics simulations to examine the structural basis for the access and binding of RvD1 to its target receptor from aqueous and membrane environments. The results offer valuable insights into the access path, potential binding pose, and key residue interactions essential for the access and binding of RvD1 to FPR2/ALXR and may help in identifying small molecule therapeutics as a possible treatment for inflammatory disorders.

## INTRODUCTION

N-formyl peptide receptor 2 or ALX receptor (FPR2/ALX), is a G-protein-coupled receptor (GPCR), associated with host immune responses such as inflammation^1^. The receptor was first identified in 1990, and Chinese hamster ovary (CHO) cells transfected with the cDNA of FPR2 displayed specific high-affinity binding to ^3^H-LXA_4_ and functional responses to LXA_4_ ^2^. Three genes encoding for human formyl peptide receptors (FPRs) have been identified, including FPR1, FPR2, and FPR3, which are clustered on chromosome 19q13.3 ^3, 4^. FPR1 and FPR2 were found to recognize various N-formylmethionine-containing peptides and play an important role in host defense and clearance of damaged host cells. However, the function of FPR3 is mostly unknown ^4, 5^. The FPRs are part of a GPCR superfamily of Gi-coupled chemoattractant receptors that belong to the γ-subgroup of rhodopsin-like class-A GCPRs ^6^. These GCPRs have seven transmembrane helices connected by three extracellular loops (ECL1, ECL2, and ECL3) and three intracellular loops (ICL1, ICL2, and ICL3) ^5^. FPR2 is expressed in a variety of non-myeloid cells, including astrocytoma cells, epithelial cells, and hepatocytes ^7^. The receptor is speculated to have various endogenous ligands that induce distinct signaling pathways to promote or resolve inflammation in ways such as wound healing and cell proliferation and has also been linked to inflammation-related diseases such as Alzheimer’s disease, rheumatoid arthritis, and systemic amyloidosis ^7, 8^. The immense functional aspects of the receptor have made it an attractive drug target.

FPRs are known to recognize a variety of ligands, including peptides containing N-formylated methionine, which can be found in products produced by bacteria and mitochondria. In addition, it is now known that many more ligands bind to the receptor, including non-formyl peptides, synthetic small molecules, and eicosanoids ^1^. FPR2 has been shown to bind to bioactive eicosanoid lipid molecules such as lipoxin A4 (LXA4) and resolvin D1 (RvD1), which have been shown to induce the resolution of inflammation. More recent studies have challenged the idea that LXA_4_ acts as a ligand for FPR2, in part due to a lack of positive control for LXA_4_-induced responses ^9–12^. RvD1 and RvD2 are both biosynthesized from docosahexaenoic acid (DHA), and the molecules belong to a specialized group of pro-resolving lipid mediators ^13^ (SPMs) that induce the resolution of inflammation ^3^. In macrophages, RvD1 was shown to stimulate cAMP/PKA signaling in an FPR2-dependent manner ^14^. Therefore, suggesting that the synthesis of SPMs, such as RvD1 and RvD2, may relieve inflammatory pain by promoting an anti-inflammatory and pro-resolution state.

Docosahexaenoic acid (DHA), an omega-3 fatty acid obtained primarily from dietary sources, is enzymatically produced in our body in response to acute inflammation ^15^. Some of the most common anti-inflammatory drugs, such as glucocorticoids, are often ineffective treatment options due to their metabolic effects, leading to osteoporosis, hypertension, dyslipidemia, and insulin resistance/type 2 diabetes mellitus ^16^. In addition, glucocorticoids work by inhibiting cyclooxygenase (COX), which can delay the resolution process. In contrast, resolvins, RvD1 and RvD2, shorten the resolution interval ^3, 8^ and demonstrate anti-inflammatory effects without the undesirable metabolic effects. The functional aspects of these molecules are well-known relative to the mechanism by which they associate and dissociate with their receptor.

Serhan and colleagues found that mice treated with aspirin and DHA would produce resolvins during the resolution phase at concentrations higher than baseline ^15^. The proposed explanation is that aspirin therapy with DHA may lead to the conversion of DHA to resolvins ^15^. A computational study to discern the activation process for the modeled ALX/FPR2 receptor using two unique agonists, aspirin-triggered 17 (R)-epimer resolvin D1 (AT-RvD1) and N-formyl-Met-Leu-Phe-Lys (fMLFK) ^17^ revealed important structural features. Using a homology model, the active and inactive states of FPR2 revealed an important structural difference found in the position of the side chain residue R123^3..50^. In the inactive state, the side chain was turned toward the cytoplasmic region, making a hydrogen bond with the C126^3..53^ residue. On the contrary, in the active state, the side chain was displaced into the receptor core, which was defined by the location of R205^5.42^. After running twenty molecular dynamics simulations (10 for AT-RvD1 and 10 for fMLFK), there was a stable hydrogen bond formed between the carboxyl of AT-RvD1 and the guanidinium group of R205^5.42^. They concluded: 1) electrostatic interactions in the cytoplasmic region of the ALX/FPR2 receptor, mainly between R123^3.50^ and S236^ECL2^, are broken during the activation of the receptor, 2) activation involves electrostatic interactions between AT-RvD1 and R205^5.42^, W254^6.48^, and Q258^6.52^, 3) R205^5.42^ is the main residue within FPR2 that establishes hydrogen bonds with agonists, 4) W254^6.48^ is the fundamental residue in opening TMH6, acting as a lever, 5) ECL3’s K269^ECL3^ residue accelerates the ALX/FPR2 receptor activation in AT-RvD1 all simulations ^17^. The scope of the paper was mainly focused on the activation process of the receptor, as shown by the inter-residues distances listed. The information on the stability of AT-RvD1 in the binding site and the feasibility of the docked pose was not given. In the supplemental information, an RMSD plot shows that AT-RvD1 rearranges (6 angstroms) almost immediately and settles into a new position, which was not shown. The receptor chosen as the template for the homology model, C5a (PDB ID 6C1R), was an inactive receptor that featured residues placed in different orientations to the new active FPR2 structures (PDB ID 6OMM, 6LW5). For example, in contrast to the new homology structures, the K192 and K269 residues face towards the helices as opposed to both residues facing outwards towards the extracellular milieu. These subtle differences can result in substantially different docked poses.

Other computational studies focus on docking non-peptide ligands using a homology model of the receptor and have also found the importance of R205^5.42^ within their docked structures. Additional authors went on to note the unique difference between peptide and non-peptide agonists. The peptide ligand was aligned in a vertical manner, while the non-peptide ligands were oriented in a horizontal position within the binding site ^18^.

Important structural observations were recorded in the crystal structure of a potent peptide agonist WKYMVm bound to FPR2 ^4^ (PDB ID 6LW5). Specifically related to the binding mode of the peptide, essential hydrophobic interactions, including V105^3.32^, L109^3.36^, V113^3.40^, L164^4.64^, F178^ECL2^, F180^ECL2^, L198^5.35^, W254^6.48^, and F257^6.51^ when mutated to alanine, were shown to reduce the EC_50_ of WKYMVm-induced IP production by over 65-fold ^4^. The authors also note that the resolved N-terminal domain may or may not be important in ligand recognition, and its position could have been affected by the crystal packing procedure. Two residues that were found to be essential in the binding of peptides were R201^5.38^ and R205^5.42^. When either was mutated, they caused significant binding impairment and decreased functional output of the receptor. R201^5.38^ in FPR2 is a hydrophobic phenylalanine in FPR3, which is likely important for receptor selectivity. The authors conclude that ligand-binding pocket in FPR2 plays an important role in regulating receptor activation and its potential for a drug target site in further molecule design.

The other available structure published ^5^ (PDB ID 6OMM) features the same potent peptide bound to the receptor along with the accompanying intracellular coupling G_i_ protein. Both structures feature the peptide in an identical orientation. An important structural feature missing from this structure is the N-terminal domain, which further suggests that the N-terminal domain may not be essential for the binding stability of this peptide. The authors once again found that when R201^5.38^ and R205^5.42^ were mutated, these mutations significantly compromised WKYMVm-induced FPR2 activation. Molecular dynamics simulations were run with the protein-peptide complex, but most analyses presented focused on the conformational changes in the receptor as opposed to ligand stability.

These studies have given us an understanding of the important residues associated with ligand stability that can guide our hand in docking RvD1. There is a gap in the understanding of how RvD1 can access FPR2, and this information can prove useful in designing therapeutics that bind to the receptor. This study outlines three methods to reach our objective: (1) docking, which helps place the ligand in a pose that satisfies previously found important interactions, such as the carboxyl group facing the arginine binding cluster, (2) unbiased simulations, which show ligand stability in the binding site with various stable electrostatic and hydrophobic interactions previously mentioned in the two structures of FPR2, and (3) association simulations which use both aqueous starting point as well as well-established membrane starting points, which were previously uncovered by PMF calculations in a DMPC bilayer for RvD1 and RvD2 ^19^. Our objective is to first identify the major binding pose of resolvin D1, uncover the relevant protein interactions, and then determine the mechanism in which RvD1 binds to the binding site of FPR2.

## MATERIALS AND METHODS

### Structures preparation and Molecular Docking

We used the crystal structure of human FPR2 in complex with WKYMVm (PDB ID 6LW5 ^4^). During crystallization, the N-terminal residues M1-E2 of FPR2 were replaced with a thermostable apocytochrome b562RIL (bRIL), and five residues at the C-terminus were truncated to facilitate crystal packing. Additionally, a single mutation S211^5.48^L (superscript denotes residue numbering using Ballesteros Weinstein nomenclature) was introduced to further increase hydrophobic interactions with nearby residues on the external surface of the receptor for protein stability ^4^. The structure was prepared for simulations using Molecular Operating Environment ^20^ (MOE) software. Finally, the mutated residues were reverted to wild type, the bRIL modification was removed, and the peptide (WKYMVm) bound to the structure was removed. Structure preparation was done through the QuickPrep tool in MOE, which assigned protonation states and rotamers, capped the N- and C-terminal ends with acetyl (ACE) and methyl amide (NME) groups, respectively, and added missing hydrogen atoms. All titratable residues were assigned to their dominant protonation states at pH 7.4. The ligand (RvD1) was prepared as deprotonated carboxylate (its most dominant species at a pH of 7.4), and charges were assigned^19^. The ligand was parameterized using the CGENFF ^21, 22^ web server. Docking simulations of RvD1 into the FPR2 binding site were carried out using MOE’s docking module, with triangular matching as the placement method with London dG scoring. A subsequent refinement was done using induced fit and the Generalized Born Solvation Integral/Weighted Surface Area (GBVI/WSA) scoring function ^23^. A docking pose with the carboxyl group of RvD1 facing the nest of arginine residues (R201^5.38^ and R205^5.42^) was deemed the relevant pose, and subsequent unbiased simulations used this pose.

### Unbiased MD and well-tempered metadynamics simulation setup

The top-scoring docked structure of the FPR2-RvD1 complex was used for unbiased MD simulations. The system was prepared using CHARMM-GUI’s membrane builder module ^24^ with the protein orientated in the bilayer using the Orientations of Proteins in Membranes (OPM) webserver ^25^. The protein was embedded in a heterogenous lipid bilayer reflecting the native plasma membrane, composed of phosphatidylcholine (POPC), phosphatidylethanolamine (POPE), phosphatidylinositol (POPI24), cholesterol, and sphingomyelin (PSM) ^26, 27^. The asymmetric bilayer had the following lipids: ∼45 POPC, ∼37 cholesterol, 22 PSM, and 5 POPE in the upper leaflet and ∼32 POPE, ∼32 cholesterol, ∼19 POPC, ∼12 POPS, ∼12 POPI24, and ∼10 PSM in the lower leaflet. A neutralizing concentration of NaCl (0.15 M) was added to the system, and water padding of ∼22.5 Å was applied on both sides using the TIP3P water model^28^. Before production simulations, the six-step minimization, heating, and equilibration simulations were carried out using the default CHARMM-GUI input parameters. A 1000-step energy minimization process, followed by a 75 ps in NVT ensemble with decreasing restraints on dihedral angles and phosphorus atoms in the membrane, was performed. Next, a 200 ps simulation in an NPT ensemble, with decreasing restraints, followed by a 100 ps simulation in NPT, without any restraints, was performed. Systems were simulated at 310 K for an additional 200 ns each using GROMACS 5.1.2 ^29^ or the updated version, GROMACS 2021. Unbiased simulations were run with the docked structure to assess whether RvD1 was stable in the binding site for at least 500 ns.

Well-tempered metadynamics ^30^ was used to elucidate the association paths and energetics involved during the binding of RvD1 to FPR2. The simulated systems had RvD1 placed in various locations near the receptor, including within the membrane, near TMH1/TMH7 or TMH4/TMH5 helices, and in the aqueous phase above the binding pocket. PLUMED ^31^ was used to set up combinations of multiple collective variables, which include 1) the distance between the center-of-mass (COM) of the ligand and the COM of three binding site residues (S288^7.39^, R201^5.38^, and Q258^6.52^), 2) the distance between the COM of the carboxyl group and the COM of R201 guanidino group, and 3) the internal angle of RvD1 defined by three atoms (the carboxyl carbon, the middle alkyl C11 and C22 at the other end). The bias factor for all association simulations was 15, the sigma values for distances were either 0.05 or 0.1, and the angle value was 0.35. We added a restraint to the COM distance between the ligand and binding site residues of 2.5 nm to decrease the likelihood of the ligand drifting away from the protein. More than 20 association simulations were run in total using a combination of the collective variables and different starting positions for RvD1 to ultimately obtain poses similar to the unbiased simulation.

### Free energy surface (FES) analysis

For all the association simulations, free energy surfaces were generated using PLUMED ^31^. The access path was chosen and plotted using the minimum energy pathway analysis for energy landscapes (MEPSA) program ^32^.

### Trajectory analysis for H-bonds and hydrophobic contacts

H-bonds between the protein and ligand were monitored using Visual Molecular Dynamics ^33^ (VMD) and the hydrogen bond plugin tool. The distance cut-off was 3.5 Å, and the angle cut-off was 45°. Protein-ligand contacts were also calculated to assess various hydrophobic interactions using an in-house Tcl script that counted all the protein-ligand interactions with a distance cut-off of 4 Å.

## RESULTS

### The potential binding pose of RvD1 was obtained from molecular docking

Multiple X-ray crystallography and cryo-EM structures^4, 5^ of FPR2 bound to high-affinity peptides have characterized the critical binding site of FPR2, which include H102^3.29^, D106^3.33^, L109^3.36^, R201^5.38^, R205^5.42^, F206^5.43^, W254^6.48^, F257^6.51^, S288^7.39^, and F292^7.43^. Specifically, both structures of FPR2 (PDB ID 6OMM and 6LW5) revealed two essential residues involved in the binding of ligands, which include R201^5.38^ and R205^5.42^. Other critical hydrophobic residues in the binding site are V105^3.32^, V113^3.40^, L164^4.64^, F178^ECL2^, F180^ECL2^, L198^5.35^, W254^6.48^, and F257^6.51^. Another docking study used both peptide and non-peptide ligands and found that many of the larger peptide molecules adopted an upright orientation in the binding site, while the smaller non-peptide ligands adopted a horizontal orientation in the binding site ^18^. The study also indicated the importance of H102^3.29^, R201^5.38^, and F257^6.51^ in ligand binding. Resolvin D1 (RvD1) was docked into the prepared FPR2 receptor (PDB ID 6LW5 ^4^) using residues R201^5.38^, H102^3.29^, and L81^2.60^ for placement of RvD1 (See the Methods section for details).

The potential binding poses were chosen based on the docking score and manually inspected for both polar and nonpolar interactions with the critical residues shown before. One of the docked poses in which the carboxyl group of RvD1 facing towards R201^5.38^ and R205^5.42^, and the alkyl core of RvD1 placed under the N-terminal residues were used for subsequent unbiased simulations to assess the stability and residue interactions of RvD1.

### Unbiased MD simulation assessing the stability of docked pose

The FPR2-RvD1 complex obtained from the docking simulations was subjected to unbiased all-atom MD simulations for 500 ns in three replicates. Although the alkyl end of RvD1 underwent notable conformational changes, the carboxyl end of the ligand appeared mostly stable and secure in the binding site (Fig. 1A and B). To further evaluate the binding pose, we quantified the polar and nonpolar hydrophobic interactions through the entire simulation time of 500 ns. Notably, the deprotonated carboxyl group of RvD1 made stable and lasting salt-bridge interactions (bond distance < 4 Å) with the positively charged guanidino groups of R201^5.38^ and R205^5.42^ (**Fig. 1B and 1C**). In addition to these charge-charge interactions, S6 and H102 formed moderately strong H-bonds (bond distance <5 Å) with the hydroxyl groups (O1, O2, and O3) of RvD1. Although the polar interactions seem to play a critical role in RvD1 binding and stability, multiple hydrophobic residues appear to engage with the ligand and contribute to the binding. The nonpolar contacts were quantified by contact occupancy (%), which gives the fraction of the simulation time during which RvD1 is within 4 Å of a given residue. The residues with >90% contact occupancy include F5 on the N-terminal domain and F292^7.43^ from TMH7, L81^2.60^, M85, V105^3.32^, V160^4.60^, L164^4.62^, and F257. Other residues, such as D106 and V284, have had contact occupancy of 60-70 % with RvD1. These interactions have been reported previously to be essential for peptides that bind to the receptor^1, 4, 5^. In addition, the orientation of RvD1 was nearly horizontal in the binding site, as reported in a previous docking study involving a variety of small molecule ligands. This horizontal orientation was postulated to be the predominant pose for small molecules, while a more vertical pose is expected for larger peptide molecules, as evident from the two FPR2 structures bound to the high-affinity synthetic peptide, WKYMVm. As presented below, in our association simulations, RvD1 adopted both horizontal and vertical orientations, likely due to the immense flexibility attributed to its 14 rotatable bonds

**Fig. 1.**
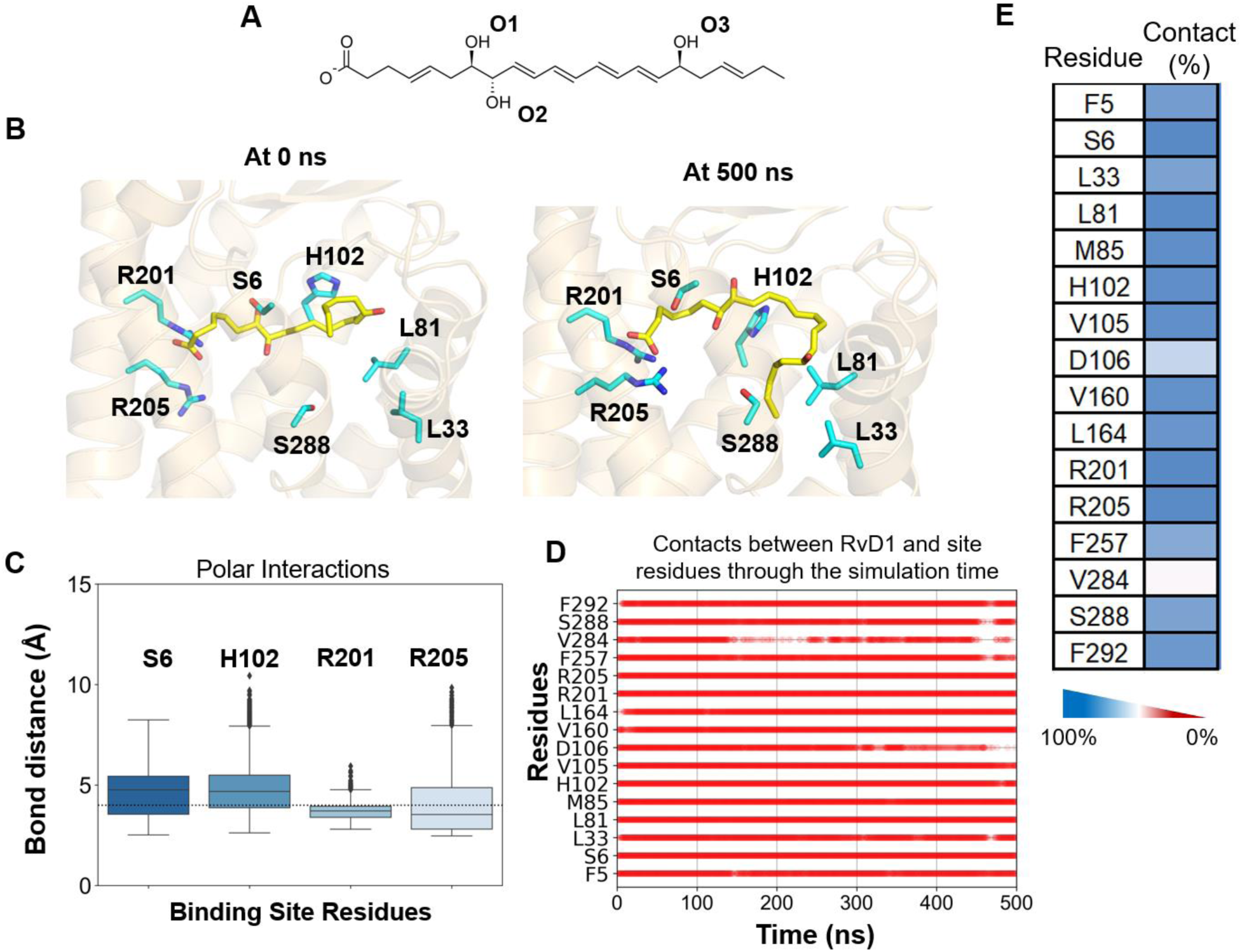
The potential binding poses of RvD1 within the FPR2 receptor site. **A)** 2D structure of RvD1 in its ionized form, the predominant species at pH 7.4. The three hydroxyl groups are labeled as O1, O2, and O3. B) The initial docked pose (0 ns) and the final pose after 500 ns of unbiased MD simulations show RvD1 forming stable H-bonds and hydrophobic interactions with the selective pocket residues. C) The distance between the polar carboxyl and various hydroxyl groups of RvD1 and the sidechain polar functional groups of S6, H102, R201, and R205 are shown. D) Contact frequency between RvD1 and the binding site residues over 500 ns simulation time. E) Contact occupancy (%) gives the fraction of the simulation time during which RvD1 was within 4 Å of the given residue.

### Association simulation results

Multiple association simulations were performed to elucidate the potential access path(s) of RvD1 to enter the binding site either from the aqueous phase above the receptor or from its energetically favorable membrane environment^19^. RvD1 was placed randomly around the receptor either within the membrane or in the aqueous phase. Out of the twenty simulations, RvD1 entered between TMH3/TMH4/TMH5 and ECL2 in five simulations. Three of the simulations demonstrated RvD1 entering between TMH5 and TMH6, while two simulations illustrated RvD1 entering directly from the aqueous phase. The collective variables used include the distance between the carboxyl group on RvD1 and R201^5.38^ and R205^5.42,^ as well as the internal angle of RvD1. Simulations were terminated as soon as the distance between the carboxyl group and arginine residues was within 4 Å. One unique characteristic of the FPR2 receptor is that the N-terminal domain sits over the binding site, which may hinder peptides and larger molecules from entering the binding site directly from the aqueous phase.

However, as RvD1 is flexible and small enough to access the binding site from various locations, the N-terminal domain on top of the binding site did not appear to hinder its entry. These initial simulations guided us in choosing other collective variables and additional starting poses for further assessments.

Additional association simulations utilized the initial simulations to inform our collective variables and assess the impact of the N-terminal domain. Simulations with a truncated N-terminal domain (residues 3-19 removed) and with a full N-terminal domain were initiated with resolvin starting on either from the extracellular aqueous phase above the receptor’s binding site or within the membrane near TMH1/TMH7, or TMH4/TMH5.

Three additional simulations of RvD1 entering from the aqueous phase were set up using two distances as the collective variable. One distance was the COM of the ligand to the binding site residues, and the other was the distance of the carboxyl group to the arginine residues, R201^5.38^ and R205^5.42^. During each simulation, RvD1 entered a pocket adjacent to the N-terminal domain, ECL2, and TMH5/TMH6 (**Fig. 3. B.)**. Each simulation ended with RvD1 in a unique conformation in the binding site, underlining the flexibility that RvD1 has. The simulations were extended for at least 100 ns in an unbiased manner after the initial entry into the binding site to assess stable conformations within the binding site. During association, RvD1 made extensive contacts with residues T177^ECL2^, F178^ECL2^, F180^ECL2^, F273^ECL3^, and L272^ECL3^ in all three simulations.

The most predominant entry path seems to be from the aqueous phase, but other association events were observed and analyzed. Figure 2 shows the important residues involved in three of the aqueous pathway simulations. RvD1 adopts a ‘vertical’ pose (**Fig. 2. D.)**, which is distinct from the unbiased simulation pose (**Fig. 1.)**. Early interactions that draw resolvin into the binding site include F180^ECL2^, F273^ECL3^, F178^ECL2^, and T177^ECL2^. Through these interactions, resolvin eventually rearranges and enters the extracellular vestibule with its carboxyl group facing R201^5.38^ and R205^5.42^. The tail portion of RvD1 drifts upward towards ECL2 and ECL3, with the carboxyl group held in place by bonds with R201^5.38^ and R205^5.42^. RvD1 orients itself within 5 Å of the unbiased simulation pose, but this pose rearranges into a vertical pose (**Fig. 3. A)**.

**Fig. 2.**
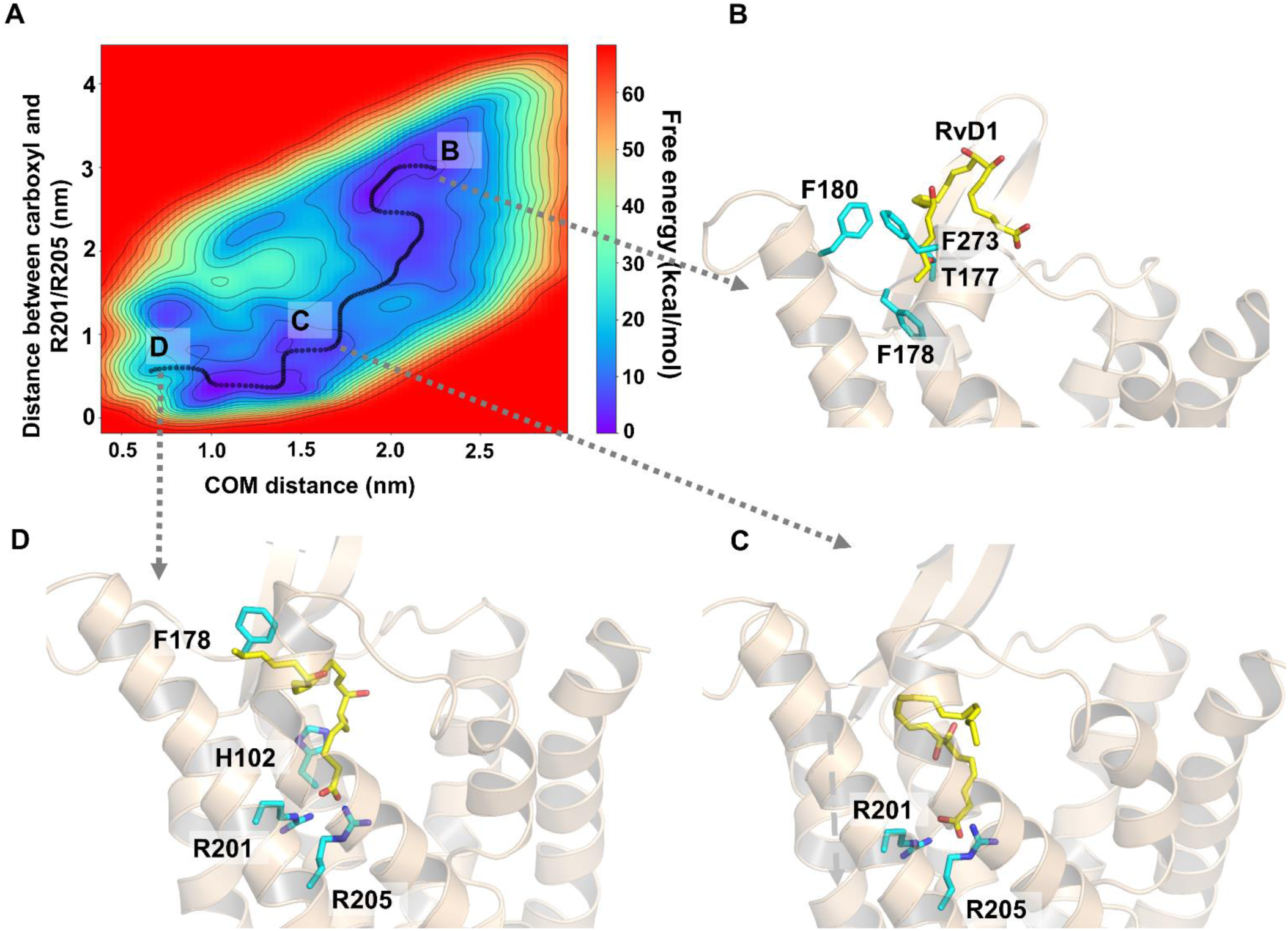
The access and binding of RvD1 into the binding site of FPR2 from the surrounding aqueous phase. A) The free energy surface shows a favorable energy profile for the association of RvD1 directly from the aqueous phase. B) Initially, RvD1 made contacts with residues F273^ECL3^, F180^ECL2^, T177^ECL2^, and F178^ECL2^ before slipping between a pocket formed by the N-terminal domain, TMH5, and ECL2. C) During association, the carboxyl group extended towards R201^5.38^ and R205^5.42^. D) Once inside the binding site, the simulation was extended for 200 ns to assess the orientation. RvD1 was flexible and adopted various orientations within the binding site. In this particular orientation, the hydrophobic alkyl tail drifts upwards towards ECL2. The N-terminal domain has been removed for the sake of clarity.

**Fig. 3.**
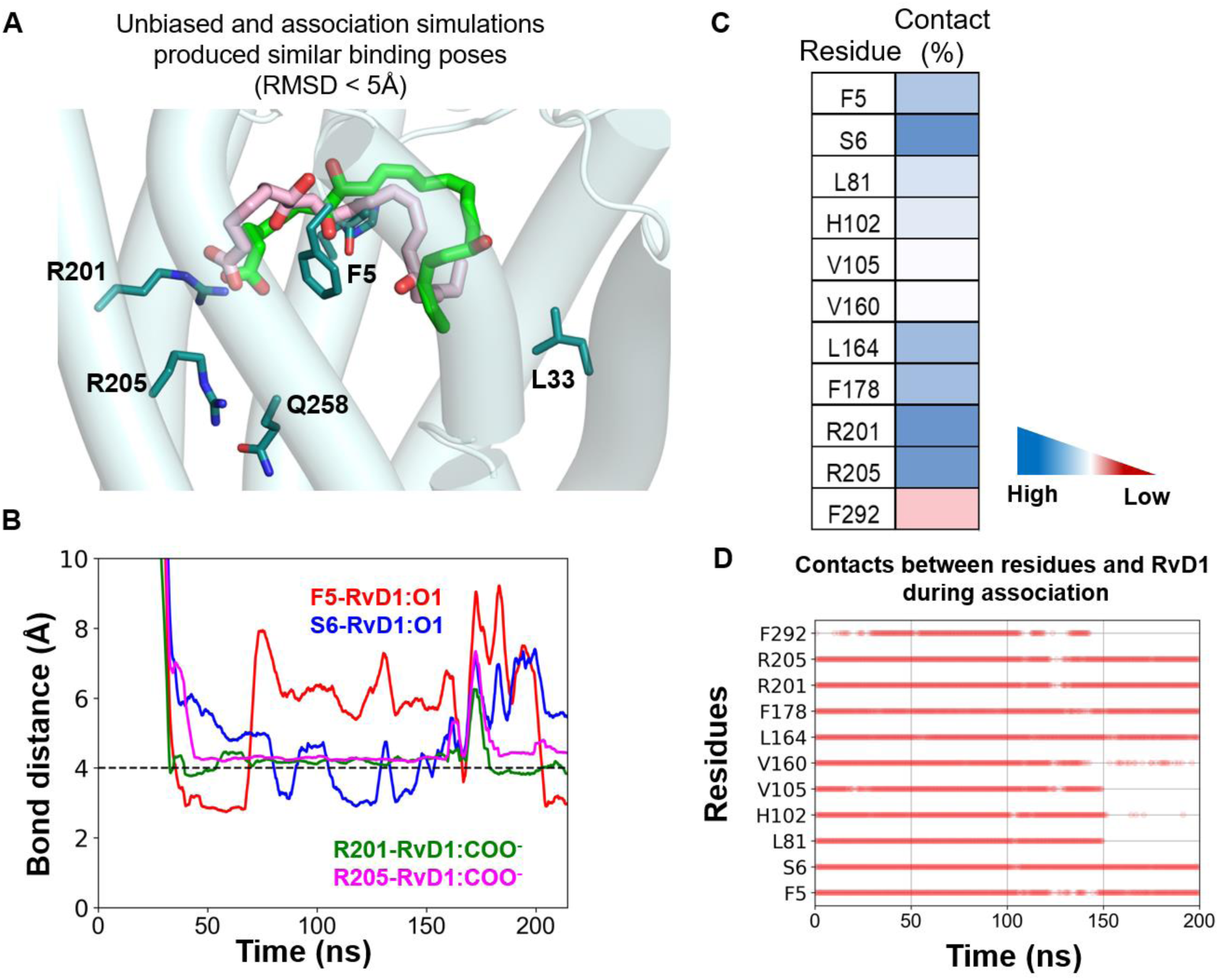
Aqueous association and extension of simulation in an unbiased manner showing important interactions. A) During the initial association from the aqueous phase, RvD1 orients itself in a similar pose to the 500 ns unbiased within 5 Å RMSD. Bias was turned off after RvD1 settled into the binding site to determine whether the binding pose would be stable, but by the end of the unbiased simulation, RvD1 adopted a vertical pose with R201^5.38^ and R205^5.42^ interacting with the carboxyl group of RvD1. A contact frequency table shows residues that were in contact with RvD1 within 4 Å over the course of the simulation B) Hydrogen bonds over the course of the simulation show important electrostatic interactions formed between R201^5.38^ and R205^5.42^ and the carboxyl group that is mostly stable through the course of the simulation. C) Contact frequency over the course of the simulation showing important hydrophobic interactions within 4 Å between RvD1 and the protein residues.

### Membrane entry through TMH5 and TMH6

Some of the simulations showed entry through TMH5 and TMH6 directly from the membrane. During this association, the carboxyl group of resolvin interacts with S211^5.48^ (**Fig. 4A)** before reconfiguring and entering the binding site with the alkyl chain entering first (**Fig. 4B)**. After the initial bias and association, the simulation was extended without any bias and RvD1 formed stable hydrogen bonds with R201^5.38^ and R205^5.42^ in an upside down ‘U’ conformation (**Fig. 4D, E)**.

**Fig. 4.**
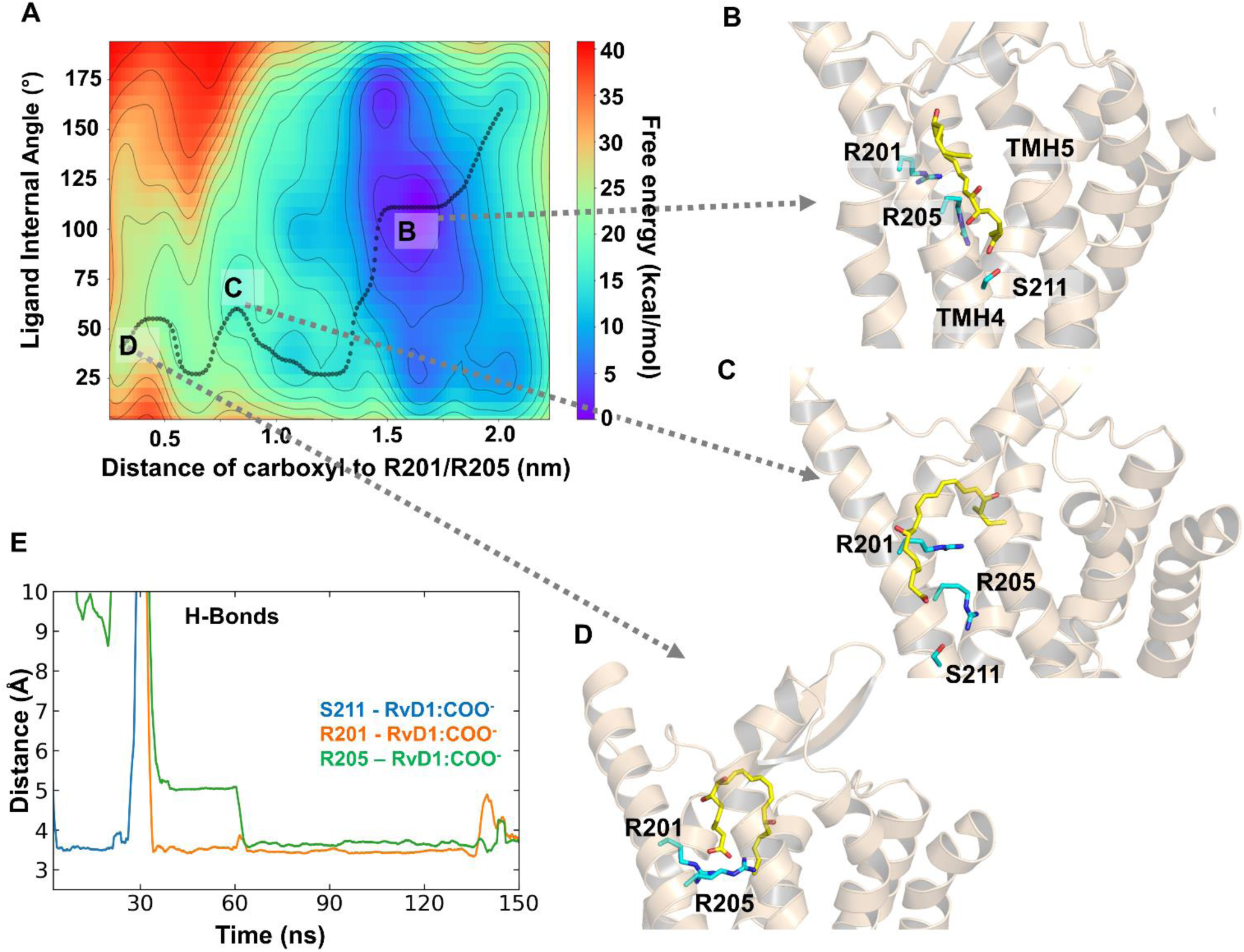
Association of resolvin D1 through TMH5 and TMH6. A) The free energy surface of the association through TMH4 and TMH5. B) Initial interactions were formed between S211^5.48^ and the carboxyl group. C) RvD1 reoriented itself, slipped between the helices, and settled into the binding pocket. D) After initial association, the simulation was extended in an unbiased manner, and RvD1 adopted an upside-down ‘U’ conformation. E) Stable hydrogen bonds were formed through association, including the important residues R201^5.38^ and R205^5.42^.

An important distinction between this membrane entry and the aqueous entry is the differences in the free energy surfaces (**Fig. 2A and Fig. 4A**). It appears that the aqueous pathway features a more energetically favorable entry than the transmembrane TMH5 and TMH6 entry. The binding pose is distinct from the unbiased simulation, but the carboxyl group is held in place by R201^5.38^ and R205^5.42^.

### Membrane entry through TMH1 and TMH2

Another plausible entry pathway for RvD1 was from the membrane and slipping between TMH1 and TMH2 before settling into the binding site in a unique extended conformation (**Fig. 5**.). The membrane entry pathway between TMH1 and TMH2 resulted in an RvD1 pose most similar to the unbiased simulation (**Fig. 1.)** but lacked the R205^5.42^ hydrogen bond featured in other simulations. This entry pathway is energetically favorable relative to the entry through TMH5 and TMH6 (**Fig. 1. A.**) and features a pose that is oriented similarly to that found in docking simulations in a horizontal manner across the binding site.

**Fig. 5.**
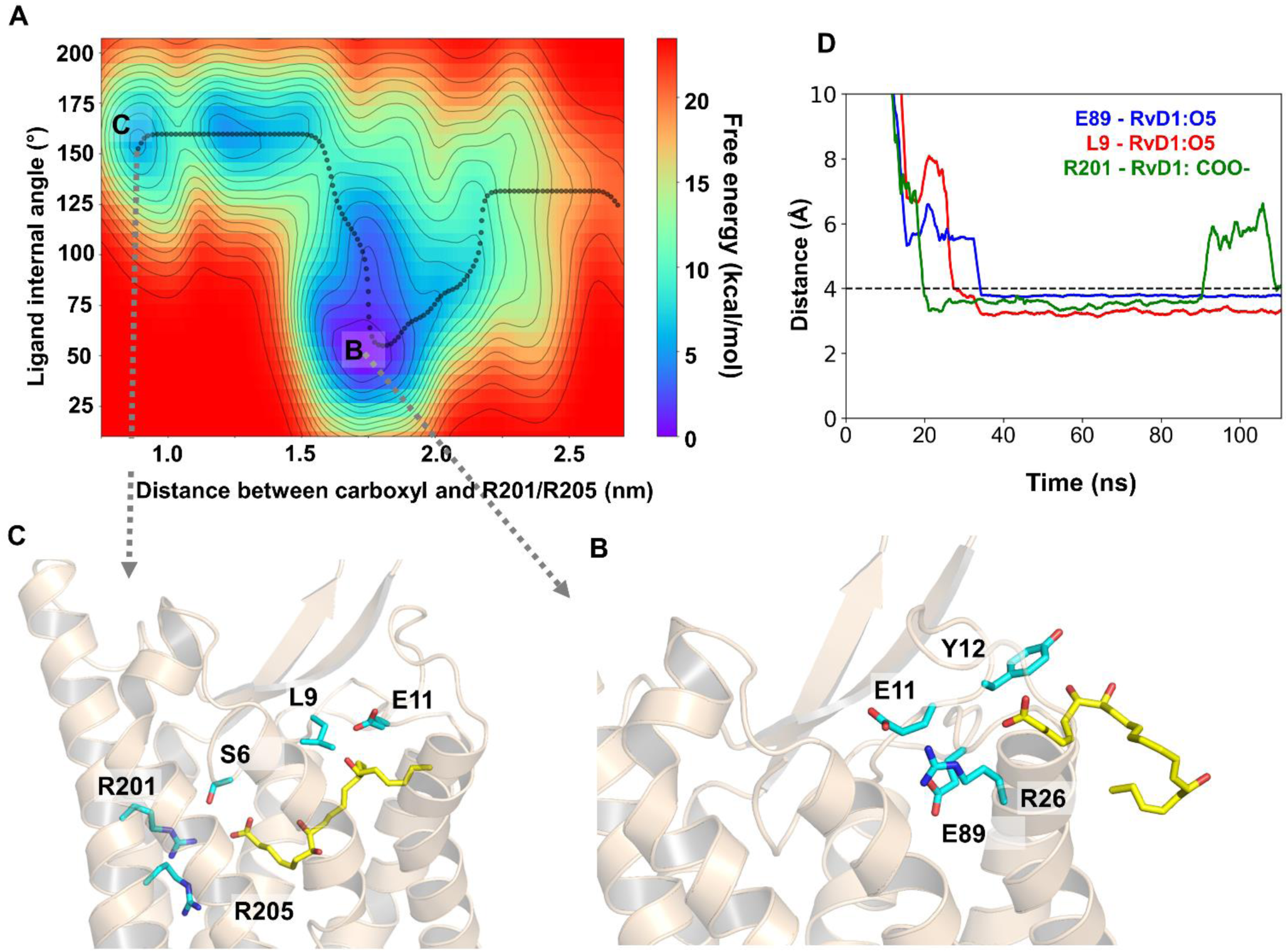
Association of RvD1 from the membrane between TMH1 and TMH2. A) The free energy surface from the membrane between TMH1 and TMH2 shows three distinct wells. B) Early in the association, RvD1 slips between TMH1 and TMH2 which involved residues L9, E11, Y12, T23^1.29^, R26^1.32^, I27^1.33^, L30^1.36^, M85^2.64^, and E89^ECL1^. C) RvD1 continues into the binding site quickly, with the carboxyl group seeking out R201^5.38^ and R205^5.42^. D) RvD1 extends across the entire binding site. The simulation was continued unbiased for 100 ns, and RvD1 was stable in the extended conformation. E) Hydrogen bonds between the hydroxyl group on the alkyl tail and E89^ECL1^ and the mainchain nitrogen of L9 held RvD1 in the binding pocket.

**Fig. 6.**
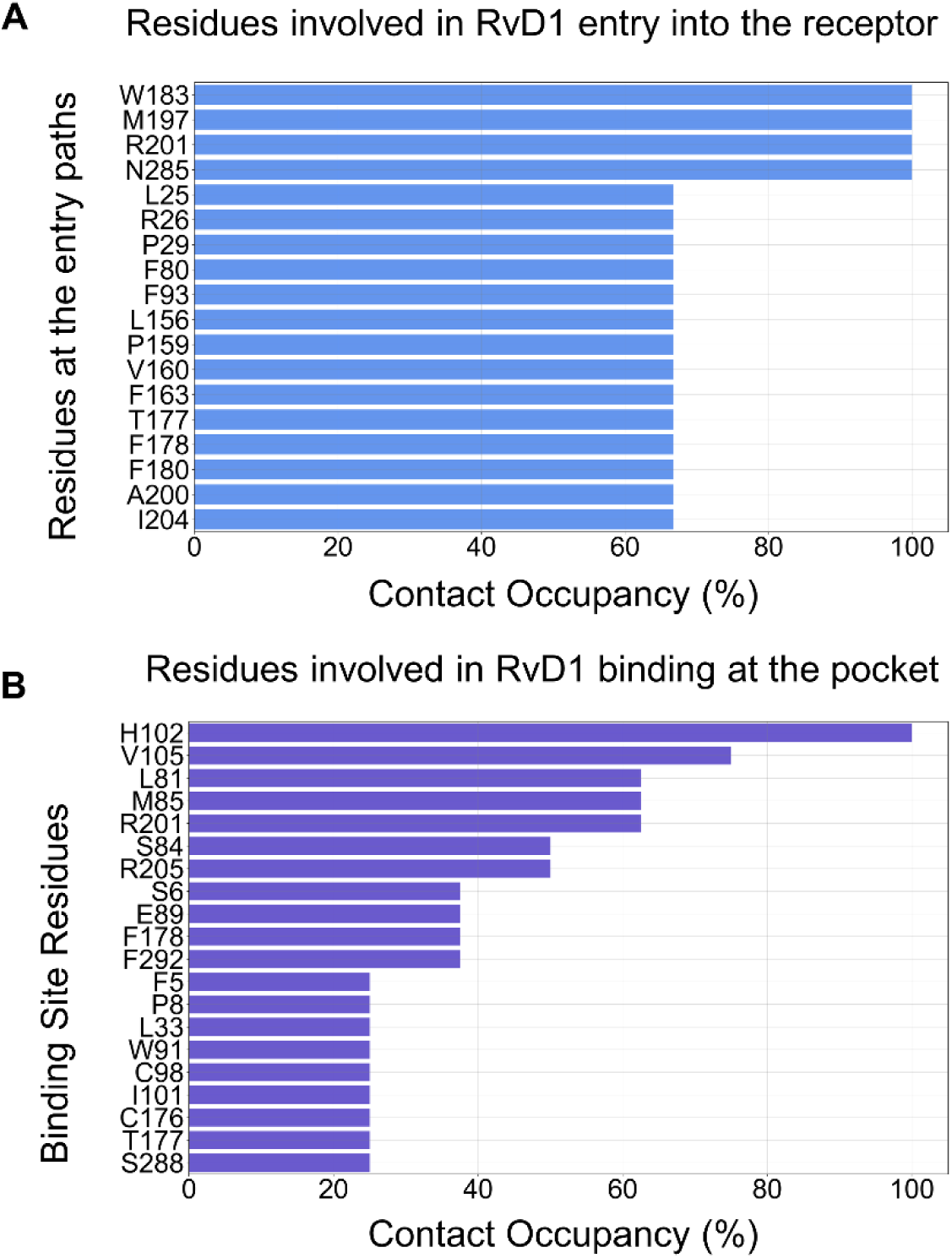
Relative contact occupancy (%) of residues interacting with RvD1 during access and binding processes observed in all the association simulations. Of the eleven simulations, contact analysis and hydrogen bond analysis were performed using VMD. Within 4 angstroms, the program records relevant interactions. These recordings were sorted from most frequently to least frequently occurring. In order to demonstrate the relative frequency of recordings, the relative ratio of the most frequently recorded residue is shown as 100%. A) R201^5.38^, M197^5.34^, N285^7.36^, W183^ECL2^, and F178^ECL2^ were the most frequently recorded residues upon entry into FPR2/ALX. B) H102^3.29^, V105^3.32^, R201^5.38^, M85^2.64^, and L81^2.60^ were the most frequently recorded resides found within the binding site. C) H102^3.29^, R201^5.38^, V105^3.32^, M85^2.64^, and R205^5.42^ were the most frequently recorded residues found collectively between both the entry pathways and the binding pocket. While H102^3.29^ has a relative ratio of 100% in C), it should be highlighted that H102^3.29^ had a relative ratio of 0% within the entry residues, meaning that it was not recorded at least once by the program throughout all eleven simulations. However, because it was recorded such a large number of times within the binding pocket, it shows up as one of the most collectively recorded residues in C).

## DISCUSSION

Resolvins represent a class of specialized pro-resolving mediators derived from omega-3 fatty acids and have been shown to bind to various G protein-coupled receptors (GPCRs), helping to resolve inflammation and restore tissue to homeostasis. Multiple classes of resolvins exist primarily based on their straight-chain polyunsaturated fatty acids or specific aspects of their structure. Resolvin D and resolvin E represent the major classes of resolvins and are further broken down into RvD1-6 and RvE1-4, which have been discovered and characterized by their unique number, position, and chirality of hydroxyl functional groups. It is known that RvD1 acts through FPR2 as well as GPR32. Due to their unique properties in downregulating a variety of inflammatory mediators and cytokines, they have emerged as a class of promising compounds to further optimize for the purposes of treating various inflammatory diseases.

In this study, twenty simulations with RvD1 starting in unique positions relative to FPR2/ALX were performed. The simulations began utilizing WT-MD (Well-tempered metadynamics) biased forces to pull RvD1 towards FPR2/ALX. Simulations using unbiased parameters were restricted by the amount of time and storage space available to perform them. Once RvD1 entered the binding site, the WT-MD biased forces were shut off, transitioning the system into an unbiased one. Three residues (H102, R205, F292) formed a triangular shape within the binding site and were used within the WT-MD system parameters to determine when the transition would take place, which occurred once RvD1 was within 4 angstroms of the three residues.

Thirty percent of the twenty simulations were evaluated to include RvD1 entering through helices III, IV, V, and ECL2. The dominant entry pathway formed a triangular shape with the helices crossing each other and RvD1 entering FPR2/ALX within the upper portion. It should be noted that not all of the starting locations for this entry pathway had RvD1 starting near helix III. The second most common entry point was between helices V and VI, with most of the starting locations for RvD1 being near helix V. While III, IV, V, and ECL2 appear to be the predominant entrance point for RvD1, it is important to consider that the number of starting locations near helix III was more than helix V, which is on the other side of FPR2/ALX. For RvD1, starting locations near helix V and between helices V and VI appear to be the most probable locations of entry.

Of the twenty simulations, nine were not further analyzed. Given that RvD1 will enter from the aqueous pathway in vivo, it was reasoned that simulations with RvD1 starting near the lower leaflet and intracellular space would have lower probabilities of entry compared to the upper leaflet and aqueous space. Of the eleven considered simulations, the entry point near the upper leaflet of helix V was still the predominant entry pathway. The upper leaflets of helices I, II, III, VII, and ECL2 were also relatively frequent entry points.

We employed various molecular dynamics to understand how RvD1 gains access to its target receptor, the FPR2 receptor. Previous MD simulations and potential of mean force (PMF) calculations showed that RvD1 prefers to embed itself in the membrane at various depths dependent on the protonation state of the molecule. We utilized the predominant deprotonated form at a pH of 7.4 and the full FPR2 structure to assess specific atomistic details of the association of RvD1. Initially, we conducted an unbiased simulation with RvD1 in the likely docked pose utilizing previous docking studies as a template for the placement of RvD1 in the binding site. The importance of R201^5.38^ and R205^5.42^ has been established as essential for both the function and affinity of ligands, and the docking process oriented the carboxyl group of RvD1 towards these residues ^4, 5^. Because RvD1 was found to localize to the membrane, we began multiple simulations from the membrane bilayer as well as multiple simulations beginning from the aqueous phase. The predominant pathway taken by RvD1 was found to be the aqueous pathway, but multiple membrane-assisted pathways were also observed. The free energy surface of the aqueous entry showed a much more favorable profile relative to the membrane entry pathways that showed energy barriers between low-energy regions. During the aqueous association, RvD1 adopts a similar pose (5 Å RMSD) to the 500 ns unbiased simulation before rearranging and adopting a vertical pose with the carboxyl group forming bonds with both R201^5.38^ and R205^5.42^. Most of our association simulations did not reproduce the unbiased pose, underlining the flexibility that RvD1 possesses even once in the binding site. Because there are no structures of RvD1 bound to a receptor, we utilized previously published information and important residues to first dock RvD1 into the binding site, but our pose may be incorrect. Association simulations revealed multiple stable poses, whereby hydrogen bonds anchored the carboxyl group with R201^5.38^ and R205^5.420^, and the alkyl tail was facing toward the extracellular side in a ‘vertical’ orientation.

## CONCLUSIONS

In this study, we performed docking, classical MD, and enhanced simulation techniques to elucidate the potential binding mode and critical residue interactions of RvD1 to its target GPCR, FPR2/ALX. Our simulations indicate that the binding site residue R201^5.38^ plays a significant role in both the entry and binding of RvD1. H102^3.29^ may have a larger role in binding, but it appears to have no significant influence on the access of RvD1 to the binding site. Resolvin D1 can take multiple access pathways into the FPR2 receptor. Despite favorably partitioning into the membrane headgroups, RvD1 seems to access the receptor through aqueous paths, in which it slides through the N-terminal end. Although membrane-mediated pathways were observed in multiple association simulations, including the entry path between TMH1 and TMH2 and between TMH5 and TMH6, the access paths through the membrane appeared to be energetically unfavorable relative to the aqueous pathways. Future studies will focus on establishing a pharmacophore model to virtually screen compounds with high affinity for the receptor and may be therapeutically useful in resolving inflammation.

## Author Contributions

BM, CTS, and PO designed and performed research, analyzed data, and wrote the manuscript. SN conceptualized and designed experiments, analyzed data, and wrote the manuscript.

## Data and software Availability

The input, parameter, and trajectory files for the unbiased simulations and association simulations through the aqueous and membrane paths can be found here. https://zenodo.org/uploads/10945623

## References

1. He, H.-Q.; Ye, R. D., The Formyl Peptide Receptors: Diversity of Ligands and Mechanism for Recognition. Molecules 2017, 22, 455.

2. Fiore, S.; Maddox, J. F.; Perez, H. D.; Serhan, C. N., Identification of a human cDNA encoding a functional high affinity lipoxin A4 receptor. J Exp Med 1994, 180, 253–260.

3. Corminboeuf, O.; Leroy, X., FPR2/ALXR Agonists and the Resolution of Inflammation. J Med Chem 2015, 58, 537–559.

4. Chen, T.; Xiong, M.; Zong, X.; Ge, Y.; Zhang, H.; Wang, M.; Won Han, G.; Yi, C.; Ma, L.; Ye, R. D.; Xu, Y.; Zhao, Q.; Wu, B., Structural basis of ligand binding modes at the human formyl peptide receptor 2. Nat Commun 2020, 11, 1208.

5. Zhuang, Y.; Liu, H.; Edward Zhou, X.; Kumar Verma, R.; de Waal, P. W.; Jang, W.; Xu, T. H.; Wang, L.; Meng, X.; Zhao, G.; Kang, Y.; Melcher, K.; Fan, H.; Lambert, N. A.; Eric Xu, H.; Zhang, C., Structure of formylpeptide receptor 2-G(i) complex reveals insights into ligand recognition and signaling. Nat Commun 2020, 11, 885.

6. Wang, L.; Yao, D.; Deepak, R.; Liu, H.; Xiao, Q.; Fan, H.; Gong, W.; Wei, Z.; Zhang, C., Structures of the Human PGD(2) Receptor CRTH2 Reveal Novel Mechanisms for Ligand Recognition. Molecular cell 2018, 72, 48–59.e4.

7. Yu, Y.; Ye, R. D., Microglial Aβ receptors in Alzheimer’s disease. Cell Mol Neurobiol 2015, 35, 71–83.

8. Bannenberg, G. L.; Chiang, N.; Ariel, A.; Arita, M.; Tjonahen, E.; Gotlinger, K. H.; Hong, S.; Serhan, C. N., Molecular circuits of resolution: formation and actions of resolvins and protectins. J Immunol (Baltimore, Md.: 1950) 2005, 174, 4345–55.

9. Hanson, J.; Ferreirós, N.; Pirotte, B.; Geisslinger, G.; Offermanns, S., Heterologously expressed formyl peptide receptor 2 (FPR2/ALX) does not respond to lipoxin A₄. Biochem Pharmacol 2013, 85, 1795–802.

10. Planagumà, A.; Domenech, T.; Jover, I.; Ramos, I.; Sentellas, S.; Malhotra, R.; Miralpeix, M., Lack of activity of 15-epi-lipoxin A₄ on FPR2/ALX and CysLT1 receptors in interleukin-8-driven human neutrophil function. Clin Exp Immunol 2013, 173, 298–309.

11. Forsman, H.; Önnheim, K.; Andreasson, E.; Dahlgren, C., What formyl peptide receptors, if any, are triggered by compound 43 and lipoxin A4? Scand J Immunol 2011, 74, 227–234.

12. Forsman, H.; Dahlgren, C., Lipoxin A(4) metabolites/analogues from two commercial sources have no effects on TNF-alpha-mediated priming or activation through the neutrophil formyl peptide receptors. Scand J Immunol 2009, 70, 396–402.

13. Serhan, C. N.; Levy, B. D., Resolvins in inflammation: emergence of the pro-resolving superfamily of mediators. J Clin Invest 2018, 128, 2657–2669.

14. Lee, H.-N.; Surh, Y.-J., Resolvin D1-mediated NOX2 inactivation rescues macrophages undertaking efferocytosis from oxidative stress-induced apoptosis. Biochem Pharmacol 2013, 86, 759–769.

15. Serhan, C. N.; Hong, S.; Gronert, K.; Colgan, S. P.; Devchand, P. R.; Mirick, G.; Moussignac, R.-L., Resolvins: a family of bioactive products of omega-3 fatty acid transformation circuits initiated by aspirin treatment that counter proinflammation signals. J Exp Med 2002, 196, 1025–1037.

16. Serhan, C. N.; Chiang, N.; Van Dyke, T. E., Resolving inflammation: dual anti-inflammatory and pro-resolution lipid mediators. Nat Rev Immunol 2008, 8, 349–61.

17. Schmitz Nunes, V.; Rogério, A. P.; Abrahão, O., Insights into the Activation Mechanism of the ALX/FPR2 Receptor. J Phys Chem Lett 2020, 11, 8952–8957.

18. Stepniewski, T. M.; Filipek, S., Non-peptide ligand binding to the formyl peptide receptor FPR2--A comparison to peptide ligand binding modes. Bio Med Chem 2015, 23, 4072–81.

19. Gc, J. B.; Szlenk, C. T.; Gao, J.; Dong, X.; Wang, Z.; Natesan, S., Molecular Dynamics Simulations Provide Insight into the Loading Efficiency of Proresolving Lipid Mediators Resolvin D1 and D2 in Cell Membrane-Derived Nanovesicles. Mol Pharm 2020, 17, 2155–2164.

20. ULC, C. C. G. Molecular Operating Environment (MOE), Montreal, QC, Canada, 2021.

21. Vanommeslaeghe, K.; MacKerell, A. D., Automation of the CHARMM General Force Field (CGenFF) I: Bond Perception and Atom Typing. J Chem Inf Model 2012, 52, 3144–3154.

22. Vanommeslaeghe, K.; Raman, E. P.; MacKerell, A. D., Automation of the CHARMM General Force Field (CGenFF) II: Assignment of Bonded Parameters and Partial Atomic Charges. J Chem Inf Model 2012, 52, 3155–3168.

23. Labute, P., The generalized Born/volume integral implicit solvent model: estimation of the free energy of hydration using London dispersion instead of atomic surface area. J Comput Chem 2008, 29, 1693–8.

24. Lee, J.; Patel, D. S.; Ståhle, J.; Park, S.-J.; Kern, N. R.; Kim, S.; Lee, J.; Cheng, X.; Valvano, M. A.; Holst, O.; Knirel, Y. A.; Qi, Y.; Jo, S.; Klauda, J. B.; Widmalm, G.; Im, W., CHARMM-GUI Membrane Builder for Complex Biological Membrane Simulations with Glycolipids and Lipoglycans. J Chem Theory Comput 2019, 15, 775–786.

25. Lomize, M. A.; Pogozheva, I. D.; Joo, H.; Mosberg, H. I.; Lomize, A. L., OPM database and PPM web server: resources for positioning of proteins in membranes. Nucleic Acids Res 2012, 40, D370–D376.

26. Ingólfsson, H. I.; Carpenter, T. S.; Bhatia, H.; Bremer, P. T.; Marrink, S. J.; Lightstone, F. C., Computational Lipidomics of the Neuronal Plasma Membrane. Biophys J 2017, 113, 2271–2280.

27. Lorent, J. H.; Levental, K. R.; Ganesan, L.; Rivera-Longsworth, G.; Sezgin, E.; Doktorova, M.; Lyman, E.; Levental, I., Plasma membranes are asymmetric in lipid unsaturation, packing and protein shape. Nat Chem Biol 2020, 16, 644–652.

28. Jorgensen, W. L.; Chandrasekhar, J.; Madura, J. D.; Impey, R. W.; Klein, M. L., Comparison of simple potential functions for simulating liquid water. J Chem Phys 1983, 79, 926–935.

29. Berendsen, H. J. C.; van der Spoel, D.; van Drunen, R., GROMACS: A message-passing parallel molecular dynamics implementation. Comput Phys Commun 1995, 91, 43–56.

30. Barducci, A.; Bussi, G.; Parrinello, M., Well-Tempered Metadynamics: A Smoothly Converging and Tunable Free-Energy Method. Phys Rev Lett 2008, 100, 020603.

31. Tribello, G. A.; Bonomi, M.; Branduardi, D.; Camilloni, C.; Bussi, G., PLUMED 2: New feathers for an old bird. Comput Phys Commun 2014, 185, 604–613.

32. Marcos-Alcalde, I.; Setoain, J.; Mendieta-Moreno, J. I.; Mendieta, J.; Gómez-Puertas, P., MEPSA: minimum energy pathway analysis for energy landscapes. Bioinformatics 2015, 31, 3853–3855.

33. Humphrey, W.; Dalke, A.; Schulten, K., VMD: visual molecular dynamics. J Mol Graphics 1996, 14, 33–8, 27-8.

